# In Vitro Reconstitution of Dynein Force Exertion in Bulk Cytoplasm

**DOI:** 10.1101/2020.05.06.081364

**Authors:** Héliciane Palenzuela, Benjamin Lacroix, Jérémy Sallé, Katsuhiko Minami, Tomohiro Shima, Antoine Jegou, Guillaume Romet-Lemonne, Nicolas Minc

## Abstract

The forces generated by Microtubules (MTs) and their associated motors orchestrate essential cellular processes ranging from vesicular trafficking to centrosome positioning [1, 2]. To date, most studies have focused on force exertion from motors anchored on a static surface, such as the cell cortex *in vivo* or glass surfaces *in vitro* [2–4]. However, motors also transport large cargos and endomembrane networks, whose hydrodynamic interactions with the viscous cytoplasm should generate sizable forces in bulk. Such forces may contribute to MT aster centration, organization and orientation [5–14], but have yet to be evidenced and studied in a minimal reconstituted system. By developing a bulk motility assay, based on stabilized MTs and dynein-coated beads freely floating in a viscous medium away from any surface, we demonstrate that the motion of a cargo exerts a pulling force on the MT and propels it in opposite direction. Quantification of resulting MT movements for different motors, motor velocities, over a range of cargo size and medium viscosities, shows that the efficiency of this mechanism is primarily determined by cargo size and MT length. Forces exerted by cargos are additive, allowing us to recapitulate tug-of-war situations, or bi-dimensional motions of minimal asters. These data also reveal unappreciated effects of the nature of viscous crowders and hydrodynamic interactions between cargos and MTs, likely relevant to understand this mode of force exertion in living cells. This study places endomembrane transport as a significant mode of MT force exertion with far-reaching consequences for cellular organization.

## RESULTS

### A 3D motility assay to study dynein force exertion in bulk

Multiple *in vivo* studies have suggested that cytoplasmic dynein may exert pulling forces on microtubules (MTs) directly from bulk cytoplasm, without contacting the cortex [12]. This mode of force exertion may have a prevalent function in many cells, as it is thought to naturally arise from motor-driven transport of vesicles, organelles and larger endomembrane networks in the viscous cytoplasm. Such forces have for instance been predicted to contribute to the outward transport of MTs that have been released from centrosomes, potentially promoting aster expansion and organization [10, 11, 15]. They may also apply a net force to asters and cause them to move, when MTs are connected to the centrosome, with no requirement for MTs to contact the cell surface [12, 16, 17]. Because longer MTs could accumulate more cargos, such pulling forces have been proposed to increase with MT length, providing a shape-sensing design for aster centration and orientation in large eggs and early blastomeres [7–9, 18]. Simple theoretical considerations suggest that a cargo moving in bulk is akin to a micro-swimmer, generating a hydrodynamic drag force scaled to its speed, its size and cytoplasm viscosity [6, 19]. To date, however, the lack of minimal reconstitution of dynein bulk pulling has hampered deciphering the basic designs of this essential mode of MT force exertion, its general relevance and its limitations.

*In vitro* surface motility assays based on motors attached to a coverslip, have populated the cytoskeleton literature in recent years, delineating essential principles of force generation in real cells [2]. We designed a 3D bulk motility assay to study how an object transported along a MT in a viscous medium, away from any fixed anchoring point, may create forces on the MTs and cause MT movement. Taxol-stabilized and fluorescently labelled MTs were mixed with dynein-coated fluorescent beads in a viscous medium, supplemented with ATP, and flowed into a microscopy chamber (Figure 1A and STAR Methods) [20]. The motor domain of *Dictyostelium discoideum* cytoplasmic dynein was biotinylated *in vivo*, and then purified [21, 22]. The dynein molecules were bound to streptavidin beads through their biotin moiety. Prior to each assay, dynein activity was quantified by performing MT gliding assays on dynein-coated coverslips and on carpets of surface-anchored dynein-coated beads (Figures S1A-C and STAR Methods). During assay optimization, we considered using different viscous agents, and selected methylcellulose (MC), because other commonly used crowders tended to over-bundle MTs, and/or strongly affect dynein activity (Figure S1D) [23, 24]. The range of MC concentrations was determined to ensure reproducible pipetting and chamber loading, while limiting MT and bead sedimentation, and also to test the influence of medium viscosity on bulk forces (see hereafter). Given these considerations, each assay allowed to capture in general ~5 events of bead gliding during a period of ~1 hour, after which most beads had reached the minus-end of MTs, where they remained bound.

At chamber mid-height, several tens of micrometers away from either chamber coverslip surfaces, MTs that appeared horizontal and to which a dynein-coated bead had bound, were immediately imaged by time-lapse fluorescent microscopy. Remarkably, the sole motion of a dynein-coated bead 0.5μm in diameter along a MT of ~30μm in length, in a viscous medium, was sufficient to cause a concomitant marked steady MT movement in the opposite direction in the microscope frame of reference (Figure 1B and Movie S1). Tracking both bead and MT ends positions revealed that, in the majority of cases, cargo moved at near-constant speeds, causing a resultant constant MT speed; and that both motions immediately stopped once the cargo reached the MT minus-end (Figure S1E). We also noticed events in which the cargo moved, and then stopped before reaching the end of the MT. These arrests presumably reflected the detachment or stalling of motors carrying the beads. Yet, in such events, the MT also moved concomitantly and in opposite direction with the cargo, and stopped when the cargo stopped. We conclude that dynein-driven cargo motion in bulk is sufficient to pull and displace MTs.

**Figure 1:**
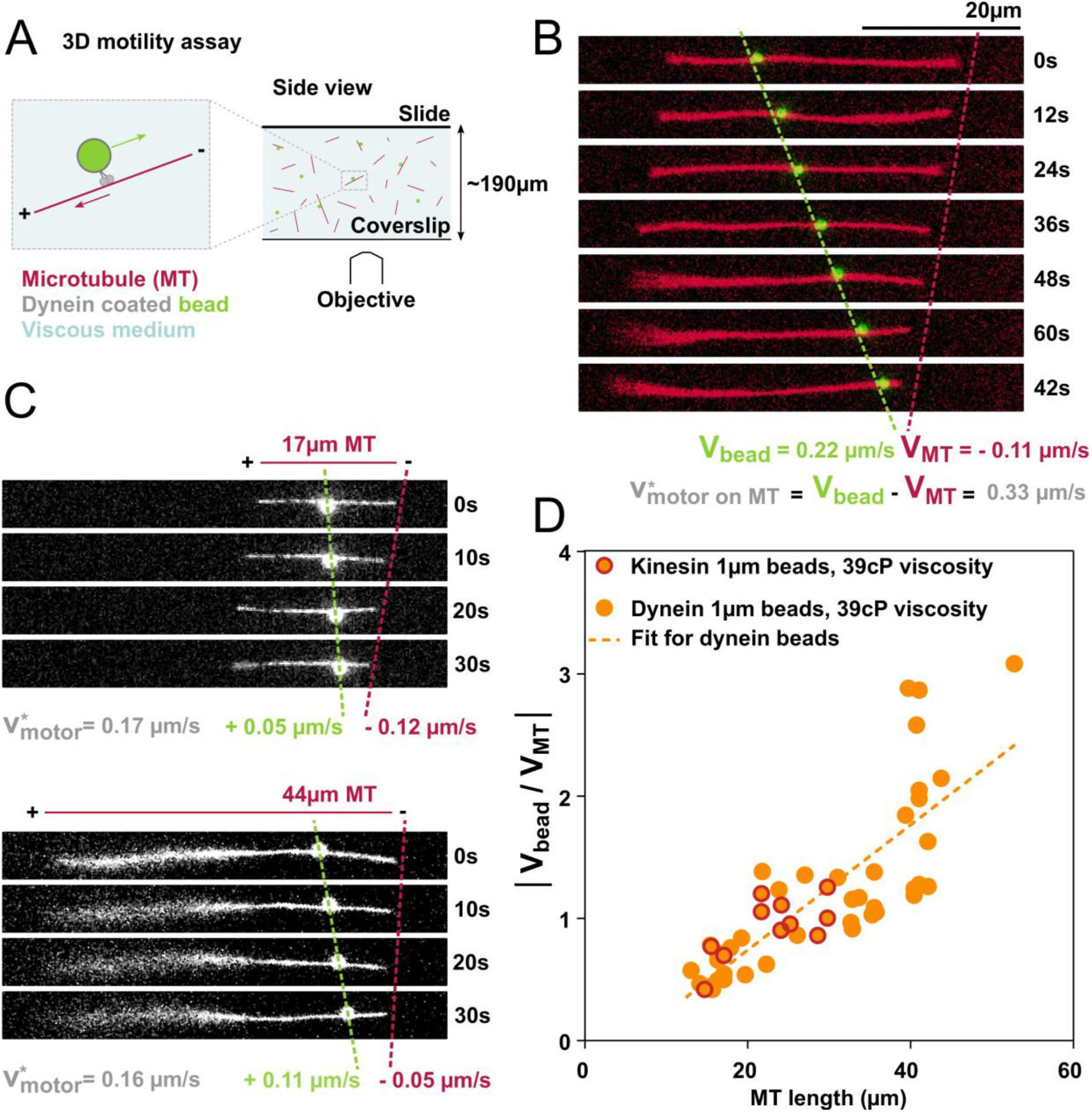
A 3D motility assay reconstituting dynein pulling on MTs in bulk. (A) Scheme representing the 3D motility assay. Taxol stabilized MTs and dynein-coated beads are mixed and flowed in a microscopy chamber, which is sealed with VALAP (see STAR Methods). MTs and beads can freely meet and the subsequent bead gliding is recorded. (B) Time-lapse of a 0.5μm diameter dynein-coated bead gliding on a MT. The motor velocity on the MT is equal to the difference between the bead velocity and the MT velocity in the lab referential. (C) Time-lapse of a 1μm diameter dynein-coated bead gliding on a short (17μm) or a long (44μm) MT. For a similar motor velocity, the shorter MT is displaced faster. (D) Bead-to-MT speed ratio as a function of the MT length, for 1 μm diameter dynein or kinesin coated beads in a 1.18% methylcellulose (MC) viscous medium (measured viscosity: 39cP). See also Figure S1.

Taxol-stabilized MTs typically ranged from 10 to 50 μm in length. We noticed that beads with the same size moving at similar speeds on a short versus a long MT, tended to displace the short MT significantly more than the long one (Figure 1C-1D and S2A). Accordingly, by computing the net velocity of the beads, V_bead_, and of the MT, V_MT_, with respect to the surrounding fluid (e.g. in the microscope frame of reference), we found that the speed ratio |V_bead_ / V_MT_|, increased with MT length in a near-linear manner (Correlation coefficient, R=0,79, R^2^ = 0.63, Figure 1D). Importantly, even though the net velocity of dynein motors along MTs exhibited some variability, likely reflecting the number of motors engaged between the bead and the MT, it did not significantly affect the dependence of the bead-to-MT speed ratio on MT length (Figure S2B-S2C). Accordingly, this behavior was also unaffected when using kinesin-coated beads, with reversed polarity and net velocity on MTs (Figure 1D and S2D). Finally, this trend was also mostly similar in single versus small MT bundles of 2-3 MTs, suggesting that MT length, and not radius was the most relevant parameter dictating MT motion in response to cargo bulk force (Figure S2E). These data are consistent with the notion that bulk MT displacement, in response to a cargo viscous force, is determined by the MT’s hydrodynamic drag coefficient that linearly scales with its length, with little influence from its diameter [2, 25].

### The efficiency of MT propulsion depends on cargo size

An important aspect of bulk hydrodynamic forces is that they are predicted to increase with the cargo drag coefficient, which increases with cargo size [25]. We thus repeated the 3D motility assay in the same conditions, but with beads of different diameters. Accordingly, for a similar range of MT lengths, we found that larger beads caused a larger concomitant MT displacement than smaller beads (Figure 2A–2B and Movie S1). These results directly demonstrate that the motion of a cargo in bulk, will drive a consequent MT displacement that respects a balance of hydrodynamic forces between the two objects, which behave as an isolated system, so that, |Y_MT_*V_MT_|= |Y_bead_*V_bead_ |, with γ the viscous drag coefficients. Importantly, because both drag coefficients are proportional to the medium viscosity, this parameter should not influence the bead-to-MT speed ratio. Accordingly, varying viscosity by changing the concentration of MC did not influence the bead-to-MT speed ratio (Figure 2C and S2F-S2G). Thus, although viscosity will linearly influence the drag forces that motors will have to overcome, it has no influence on the speed ratio between the cargo and the MT. With the ranges of cargo size, motor velocity and medium viscosity that we explored in our assays, we estimated based on Stokes formula, that net bulk forces ranging from a few hundredths to a few tenths of pN were applied to MTs. These modest forces are sufficient to move MTs of tens of micrometers at typical speeds of few tenths of μm/s. In the cell context, where several motors collectively move cargos of similar sizes at velocities of ~1 μm/s in a cytoplasm typically 100-1000x more viscous than water [26], the resulting forces are expected to be significantly higher, on the order of several pN, comparable to forces exerted by cortex-anchored motors [27, 28].

**Figure 2:**
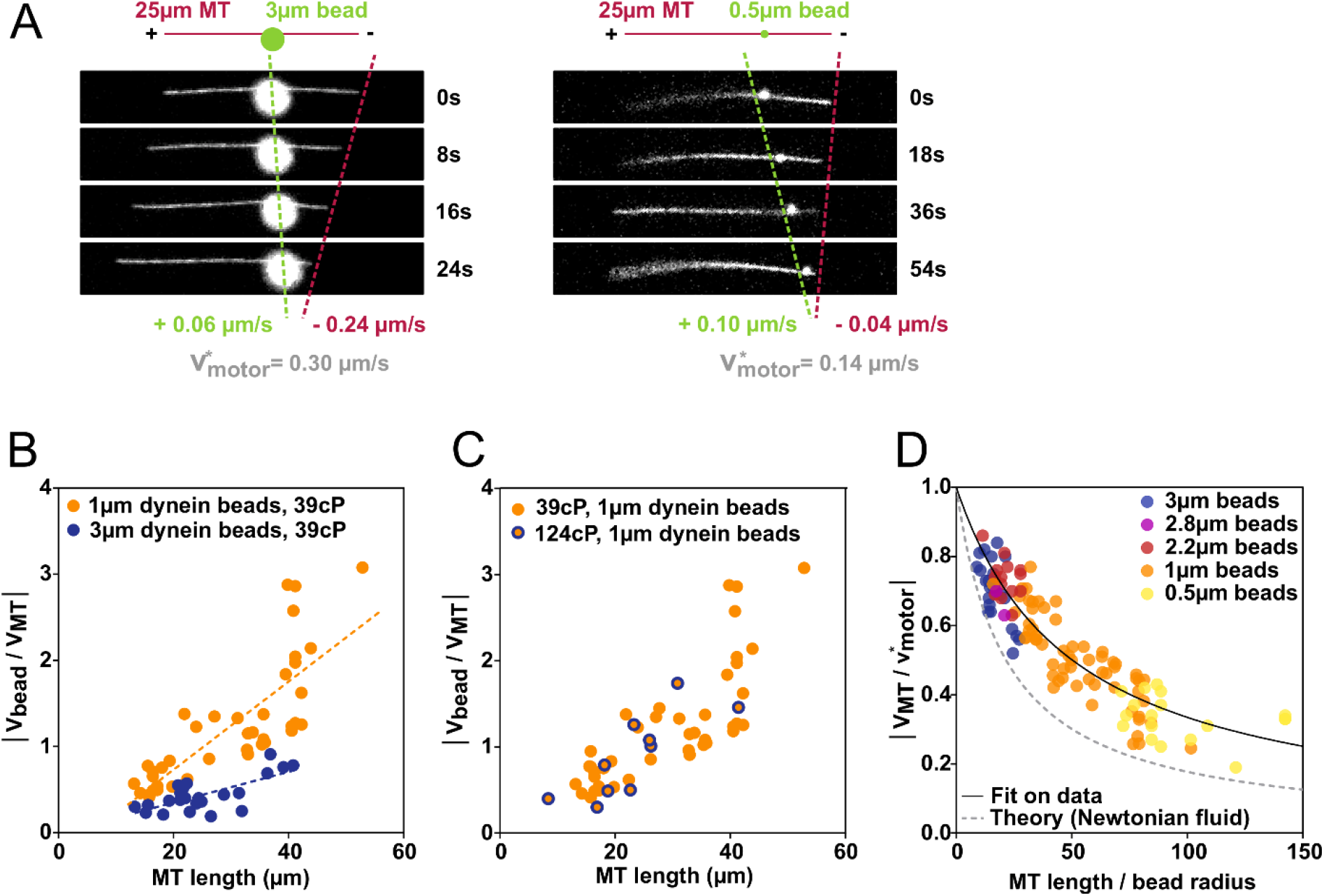
Impact of cargo size on MT bulk pulling efficiency. (A) Time-lapses of a 0.5 or 3μm diameter dynein-coated bead gliding on a ~25μm MT. For a similar MT length, the larger bead will displace the MT more than the smaller bead. (B) Bead-to-MT speed ratio plotted as a function of MT length, for dynein-coated beads of 1 μm and 3 μm diameter in a 39cP MC viscous medium. The dotted lines are linear fits to guide the eyes. (C) Bead-to-MT speed ratio plotted as a function of MT length, for 1 μm diameter dynein coated beads in a 1.18% or 1.60% MC medium (respectively yielding measured viscosities of 39cP and 124cP). (D) MT-to motor speed efficiency ratio plotted as a function of the ratio of MT length to bead radius, for beads of diameters of 0.5, 1,2.2, 2.8 and 3μm, in medium of viscosities of 39cP or 124cP. A fit of the data is shown as a black line, and the theoretical behavior in a Newtonian fluid as a grey dashed line. See also Figure S2 and S3.

Since the extent of MT propulsion appeared to depend solely on MT length and bead diameter, we next sought to determine if all our data followed a master curve, as a function of the MT-length-to-bead-radius ratio (L_MT_/R_bead_), which is a first-order approximation for the ratio of their viscous drag coefficients (Figure S3A and STAR Methods). For this, we computed an MT-to-motor speed ratio (|V_MT_/v_motor_*|), that we termed “efficiency ratio”, as it represents how much of the motor activity is converted into MT movement and is comprised between 0 and 1. A value close to 0 corresponds to a situation where the motor activity is mostly converted into bead movement, with little MT motion, much like a vesicle trafficking on a static MT network. Conversely, a value close to 1 corresponds to a large MT motion, and little bead motion, mimicking MT outward transport [10, 11, 15]. By plotting this efficiency ratio, as a function of L_MT_/R_bead_, for all bead and MT sizes, we found that our experimental conditions covered all values of efficiency (Figure 2D). To compare our data with a simple theoretical situation, we computed this curve assuming the medium was a Newtonian fluid, and neglected potential effects such as the hydrodynamic coupling between the bead and the MT, or the rotation of the bead as it moves along the MT (STAR Methods). We find that our data are systematically above this theoretical curve, but follow the same trend (Figure 2D). In fact, our data can be well fitted by the same theoretical formula, with a 2.3-fold reduction of the MT drag coefficient relative to the bead drag coefficient (Figure 2D, STAR Methods). This reduction of the effective MT drag coefficient favors its movement, and could be caused by the non-Newtonian nature of the medium, and an effect known as “shear thinning”, where the MC polymers in the medium would be locally stretched along the highly anisotropic MT shape, thereby reducing its viscous friction as it moves [29, 30]. Alternatively, this effect could also be caused by scale-dependent viscosity, when the size of the object (here the radius of the MT) approaches the mesh size of the polymer solution [31].

### Collective effects of multiple cargos pulling on MTs and minimal “asters”

To tackle another physiological situation where more than one cargo is applying bulk forces to a MT, we investigated the collective effect of several beads moving on a MT. Such experiment is relevant to aster centration, where it has been proposed that longer MTs could accumulate more cargos, thereby applying larger forces to centrosomes [8, 9, 17]. Given that, in our 3D motility assay, the number of events where two beads encounter a MT naturally was very low, we used an optical trap to place two beads on the MT (Figure 3A and Movie S2). We observed that MTs with two beads moving along them were displaced more than MTs of comparable length with only one bead (Figure 3B and Movie S3). Data points obtained with 2 beads also fell on the master curve of Figure 2D, considering the average motor speed and the sum of the bead radii (Figure 3C and STAR Methods). Importantly, we did not notice any obvious slowdown of MT speed, even when beads were moving close to each other, suggesting that putative hydrodynamic screening between cargos do not significantly affect the additivity of forces (Figure S3B). These data show that the forces exerted by multiple cargos are additive, providing an *in vitro* validation that longer MTs with more cargos may be pulled more.

**Figure 3:**
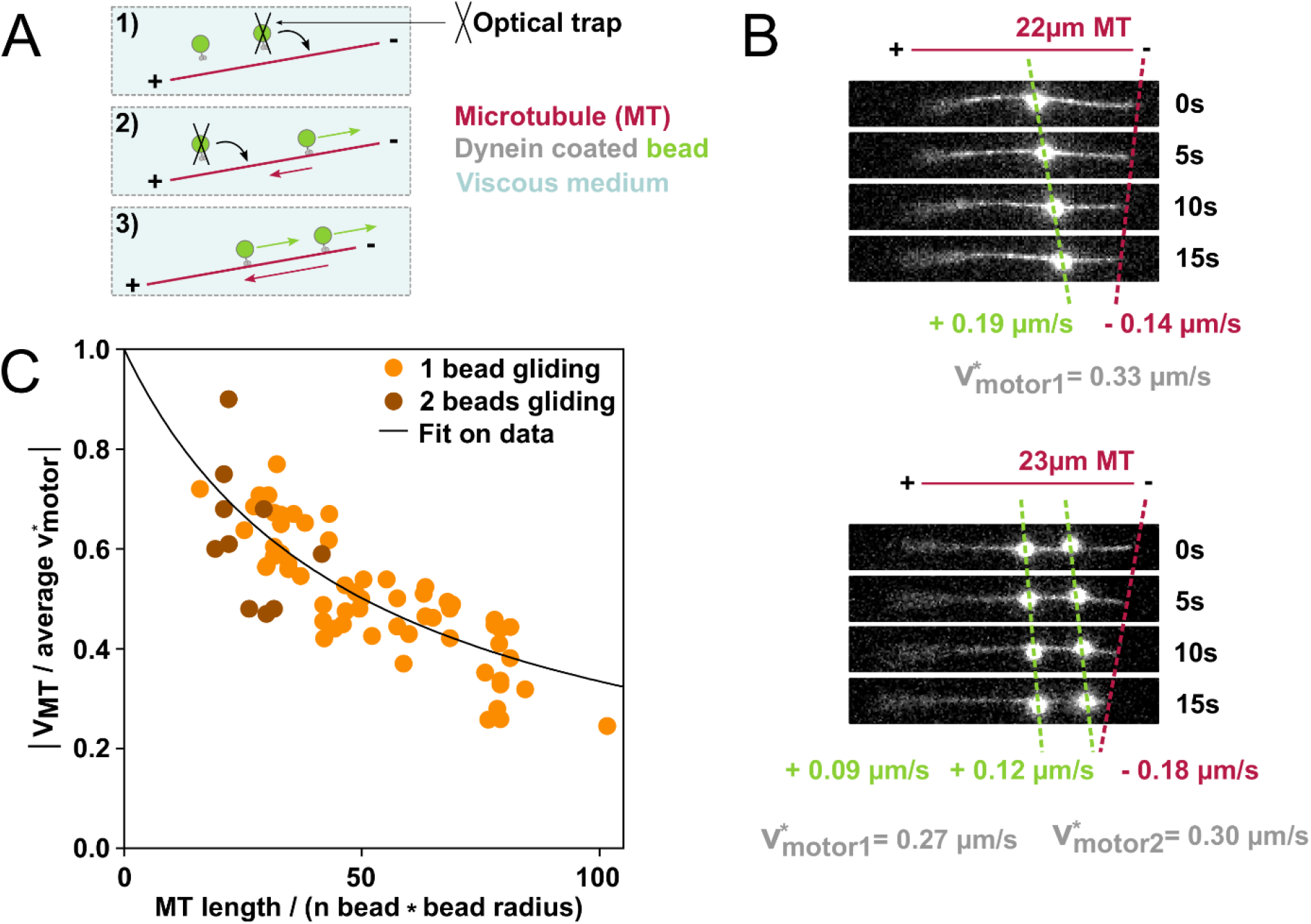
Collective effect of two beads pulling on a single MT. (A) Scheme showing how two dynein-coated beads are placed on a MT using an optical trap, to facilitate this assay. (B) Time-lapse of one or two beads walking on ~22μm long MTs. For similar motor velocities, the MT is displaced more when two beads are walking on it. (C) MT-to motor speed ratio plotted as a function of the ratio of the MT length to the bead number multiplied by the bead radius. Data shown are 1μm diameter beads, walking on MTs alone or by two, away from each other (the orange dots contain data with beads coated with dynein or kinesin, in a viscous medium of 39cP or 124cP). The fit on the data of all bead sizes is shown as a black line.

We next investigated another important situation where these forces could pull in different directions, generating a tug-of-war. To do so, we built artificial, minimal asters, consisting of two or three MTs. These minimal asters were built using dynein-coated beads and the optical trap. Briefly, we first placed a bead on a MT, let it reach and stop at the minus end, and repeated this operation with the same bead on another MT (Figure S3C and STAR Methods). This resulted in a construction made of two or three MTs, with their minus ends bound to a single bead that we refer to as the “aster center”. Using the optical trap, we then placed two or three other beads on MTs, near their plus ends, and tracked their movements as well as the concomitant movement of the aster center (Figure 4). This assay thus allows to reconstitute the basic elements of more complex *in vivo* situations, where several objects pull on several MTs, for example when organelles pull on front vs back MTs of a centering a aster [6].

Minimal asters made of two aligned, antiparallel MTs provided a “tug-of-war” situation (Figure 4A and Movie S4) illustrating that a minimal aster will move toward the region where more cargos are being transported. Following hydrodynamics force balance, the velocities of different minimal asters were here also dictated by the vectorial sum of cargo velocities (Figure 4B). To further illustrate the generalization of this principle to more complex bi-dimensional asters, we also tracked the 2D movement of a minimal aster made of three MTs and three motile beads (Figure 4C-4D and Movie S4). As above, we could detect that the movement of the aster center globally followed the 2D direction and amplitude, set by the vectorial sum of motile cargos velocities, even when the motion of beads became relatively small. We thus showed that cargos gliding on MTs of a minimal aster will displace this aster in the direction set by the vectorial sum of the hydrodynamic forces exerted on these objects, with a speed scaled to the net sum of cargos speeds.

**Figure 4:**
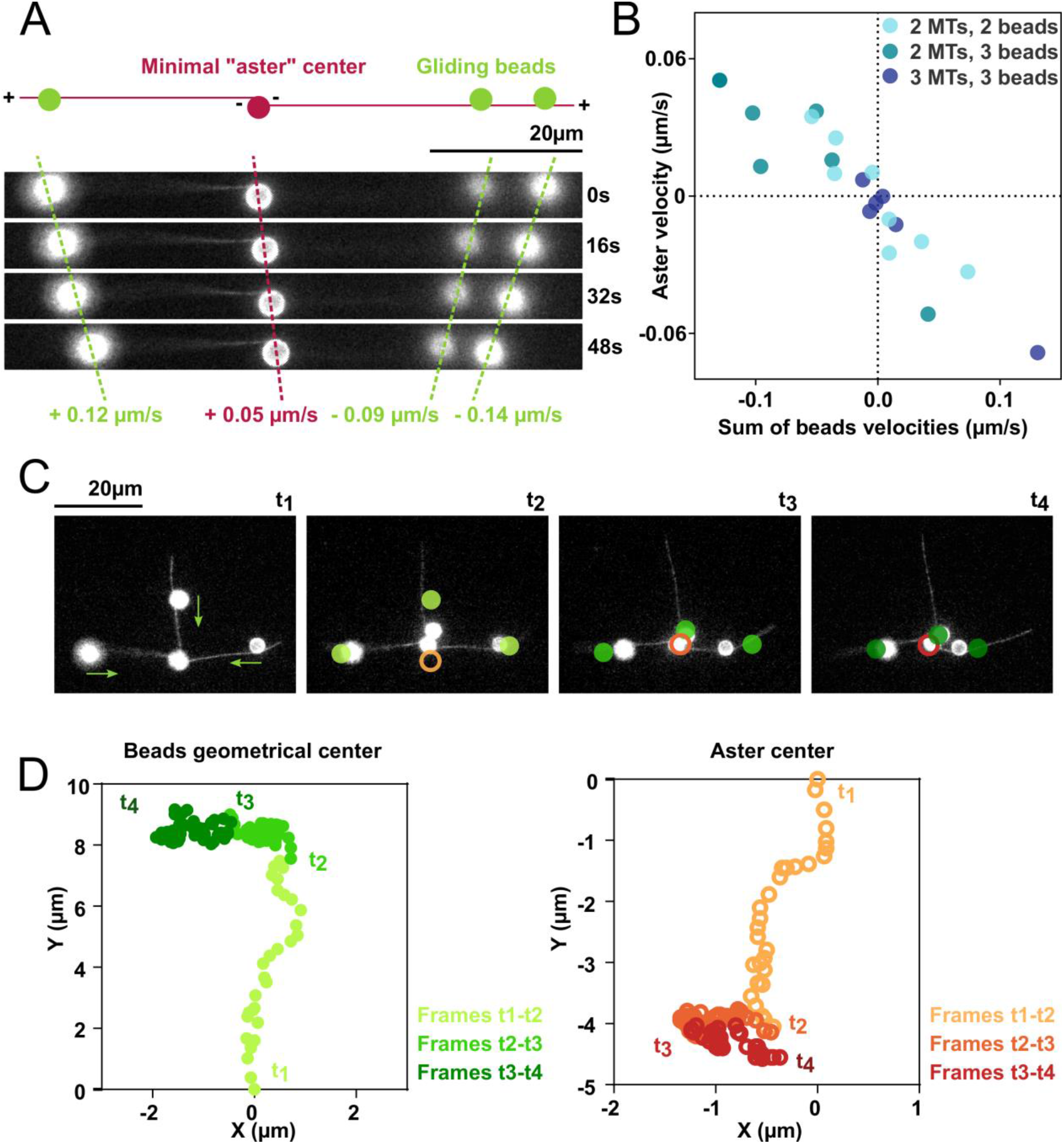
Beads pulling on MTs in bulk can move a minimal “aster” in 2D. (A) Time-lapse of three dynein-coated beads gliding on a minimal aster composed of two antiparallel MTs bound together by a dynein-coated bead at their minus ends. (B) Aster center velocity plotted as a function of the sum of dynein-coated gliding beads velocities, for tugs-of-war between two or three motile beads walking on a two-MT minimal aster, or three motile beads walking on a three-MT minimal aster. (C) Time-lapse showing three dynein-coated beads pulling on a minimal aster made of 3MTs (3μm beads, in a 39cP MC viscous medium). Walking beads at time t1 are shown as plain circles and the bead marking the “aster center” bead at time t1 is shown as a hollow circle. (D) Positions of the geometrical centers of the dynein-coated beads gliding along MTs, and of the aster center computed from the time-lapse shown in 4C. See also Figure S3.

## DISCUSSION

We here report on the development of a minimal 3D bulk motility assay to study and evidence the role of bulk motor forces on MTs. This assay shows that a mobile cargo moving along a MT exerts a force on that MT from within the viscous medium, which is sufficient to displace the MT over long distances of tens of μm. By spanning bead sizes and MT lengths, we reproduce a range of physiological behavior, from bare centripetal trafficking, to MT outward propulsion reported in many cells. Reconstituted tugs-of-war show that minimal asters will move towards the region where more cargos are being transported, essentially recapitulating proposed models *in vivo*, for aster self-propelling motion that follows the direction of asymmetries in MT lengths and/or bulk cargo densities [6, 8, 9, 17].

In our assay, cargos with all types of directionalities and sizes may effectively apply forces to move MTs. However, the important question of which specific cargos effectively promote aster centration in cells remains mostly open, although specific types of trafficking vesicles have been proposed in some systems [6, 12, 13]. Large cargos, like the endomembrane networks of the Endoplasmic Reticulum, may have higher effective drag coefficients, but also require more motors to be pulled at the same speed than small cargos. The efficiency of cargo-based bulk pulling for a dense MT aster network is also predicted to be affected by the surrounding MTs in the aster, because of the putative complex hydrodynamic interactions between cargos and neighboring MTs [32]. As an evidence for this, a simple assay using our optical trap, shows that the motion of a non-coated bead close to a free MT can move the MT with a speed up to ~20% of the bead speed (Figure S4A-S4C). In a dense aster, we speculate that this effect may become more important, plausibly limiting force transmission from cargos to MT asters. Our assay also indicates that the non-newtonian nature and shear-thinning properties of a dilute polymeric solution, or mesh-size effects, could significantly reduce the longitudinal drag coefficient of MTs, thereby facilitating their displacement over that of cargos. These results may have direct relevance to the cytoplasm which is a complex non-newtonian fluid filled with polymers and membranes, and which also exhibits shear-thinning properties, and a scale-dependent viscosity [33, 34]. Minimal *in vitro* MT force generation assays, have paved the way to our understanding of the role of MT forces in cells [3, 35]. We foresee that further complexification of our initial minimal 3D bulk motility assay, potentially incorporating complex fluids closer to the cytoplasm, cytoskeletal networks or more physiological groups of active motors, will serve to decipher the key physical and biological elements promoting and limiting bulk dynein hydrodynamic force exertion in cells.

## SUPPLEMENTAL INFORMATION

The Supplemental Information includes four figures and four videos.

## ACKNOWLEDGMENTS

We gratefully thank Carsten Janke from Curie - Orsay for providing us with porcine brain tubulin, and the mouse kinesin plasmid. We also thank Bérengère Guichard for her valuable technical help, and Dmitry Ershov and Serge Dmitrieff for their help with software and statistical analysis, and all the members of the teams “Cellular spatial organization” and “Regulation of actin assembly dynamics” for fruitful discussions. We also thank our colleagues A. Guichet, J. Heuvingh and O. Du Roure for carefully reading the manuscript. This work was supported by the Centre National de la Recherche Scientifique (CNRS), the European Research Council (ERC CoG Forcaster no. 647073) to N.M., (ERC StG BundleForce no. 679116) to A.J, and Japan Society for the Promotion of Science KAKENHI (18K06147 and 19H05379) to T.S. H.P. acknowledges the “École Doctorale FIRE - Program Bettencourt” and a fellowship from the “Fondation pour la Recherche Médicale” (FRM, FDT201904008226).

## AUTHOR CONTRIBUTIONS

Conceptualization, N.M., G.R.L. A.J. and H.P.; Methodology, T.S., J.S. and B.L. and H.P..; Investigation, N.M., G.R.L., A.J., K.M., T.S., J.S., B.L. and H.P.; Writing –Original Draft, N.M., G.R.L and H.P. Draft Editing N.M., G.R.L., A.J., K.M., T.S., J.S., B.L. and H.P.

## STAR METHODS

### CONTACT FOR REAGENT AND RESOURCE SHARING

Further information and requests for resources and reagents should be directed to and will be fulfilled by the lead contact, Nicolas Minc.

### METHOD DETAILS

#### Proteins expression and purification

##### - Dynein

Recombinant cytoplasmic dynein was expressed in *Dictyostelium discoideum*, and purified following the protocol described by T. Kon *et al*. [36] with some modifications. This HFB380 380 kDa recombinant dynein was engineered from the *Dictyostelium discoideum* cytoplasmic dynein heavy chain gene as described previously [21]. Affinity tags were added (His6 and FLAG) at the N-terminus as well as a N-terminal BioEase tag for *in vivo* biotinylation. Glutathione S-transferase (GST) could also be added for dynein dimerization. Finally, a MB35 plasmid (ID 44 - Dicty Stock Center) encoding a tetracycline-controlled transcriptional activator was introduced in order to control the recombinant protein overexpression.

Cells were cultivated at 21°C in culture dishes in HL-5 medium supplemented with 10μg/mL G418 and 10μg/mL tetracycline until they reached confluence, and were then transformed through electroporation with an MB38-based plasmid containing the engineered dynein gene. 10μg/mL blasticidin was added for plasmid selection one day after electroporation. 200mL of HL5 supplemented with G418, tetracycline and blasticidine were inoculated in a confluent 10cm culture dish, and cultivated at 21°C, 200rpm. After cells had reached 10^7^ cells/mL, dynein expression was induced through removal of tetracycline (centrifugation: 1,000g, 5min, 21°C), and resuspension in 400mL HL-5 supplemented only with G418 and 20μM of d-biotin for dynein biotinylation.

Cells were harvested by centrifugation (5,000g, 5min, 2°C), washed in 20mL lysis buffer (100 mM PIPES-KOH, 4 mM MgCl2, 0.1 mM EGTA, 0.9 M glycerol, 10 mM imidazole, pH 7.0), and resuspended in an equal volume of lysis buffer supplemented with 1 mM TCEP, 10 mM ABESF, 0.2 mM leupeptin, 87 μM pepstatin, 10 mM TAME and 0.1 mM ATP. Cells were sonicated (six pulses of 3s with 30s pause between each pulses) and the supernatant was collected after two successive centrifugations (18,000g, 20min, 2°C and 100,000g, 15min, 2°C) for clarification of the lysate. This lysate was gently mixed with 300 μL of pre equilibrated Ni-NTA agarose for 1h at 4°C on a rotating wheel. The mixture was loaded on a column, column flow though was discarded, and 12 column volumes (CV) of wash buffer (100mM PIPES, 4mM MgCl2, 0.1mM EGTA, 0.9M glycerol, 20mM imidazole, 1 mM TCEP, 10 mM ABESF, 0.2 mM leupeptin, 87 μM pepstatin, 10 mM TAME and 0.1mM ATP, pH 7) were flowed in and discarded. Bound proteins were then eluted with 2CV of elution buffer (100mM PIPES, 4mM MgCl2, 0.1mM EGTA, 0.9M glycerol, 250mM imidazole, 1 mM TCEP, 10 mM ABESF, 0.2 mM leupeptin, 87 μM pepstatin, 10 mM TAME and 0.1mM ATP, pH 7). The eluate was supplemented with 150mM NaCl, 5mM EGTA and 0.1mM EDTA, and then gently mixed with 100 μL of pre equilibrated AntiFLAG gel for two hours at 4°C on a rotating wheel. The gel was loaded on a column, column flow through was discarded, and the gel was washed with 10CV of PMEGS buffer (100mM PIPES, 4mM MgCl2, 5mM EGTA, 0.1mM EDTA, 0.9M glycerol, 200mM NaCl, 1 mM TCEP, 10 mM ABESF, 0.2 mM leupeptin, 87 μM pepstatin, 10 mM TAME and 0.1mM ATP, pH 7) and then 10CV of PMEG30 buffer (30mM PIPES, 4mM MgCl2, 5mM EGTA, 0.1mM EDTA, 0.9M glycerol, 1 mM TCEP, 10 mM ABESF, 0.2 mM leupeptin, 87 μM pepstatin, 10 mM TAME, 20% (w/v) trehalose and 0.1mM ATP, pH 7). Bound recombinant dynein was eluted slowly with 3CV of PMEG30 buffer supplemented with 0.25mg/mL FLAG peptide. The final eluate was flash frozen using liquid nitrogen, and stored at −80°C. Protein concentration was determined using A_280_ or Bradford reagent with bovine serum albumin as a standard.

##### - Kinesin

His-tagged recombinant kinesin (pET28-mKif5B_N1665, mouse kinesin N-terminal, with motor domain and coiled coil) was expressed in Rosetta (DE3)pLysS E.coli. One liter of 2YT media supplemented with 34μg/mL chloramphenicol and 50μg/mL kanamycin was grown at 37°C, 200 rpm until the OD_600nm_ was between 0.8 and 1 and transferred to 20°C. Protein expression was then induced for 4-5h at 20°C by addition of 0.5 mM IPTG.. Cells were harvested by centrifugation (8,000g, 20min, 20°C), and the pellet was resuspended in 40mL of lysis buffer (20 mM potassium phosphate, 150 mM NaCl, 10% glycerol (w/v), 5mM ß-mercaptoethanol, 1mM MgCl2, 0.1mM ATP.). Cells were then sonicated (5 pulses of 15s, 50% intensity, 6mm probe), and the lysate was clarified by centrifugation (80,000g, 50min, 4°C). The supernatant was then mixed with 0.250mL Ni-NTA beads (pre equilibrated in equilibration buffer: 50 mM potassium phosphate, 100mM NaCl, pH 7) and incubated for 1h at 4°C under gentle agitation. The resin was washed with minimum 5CV of wash buffer (50mM potassium phosphate, 10% glycerol, 2mM ß-mercaptoethanol, 25mM imidazole and 1M NaCl, pH 7) and the kinesin was eluted with 1CV of elution buffer (150mM imidazole, 50mM KCl, 10% glycerol and 10mM ß-mercaptoethanol, pH 7). The eluate was dialyzed against 50 mM MOPS, 250mM NaCl, 5mM ß-mercaptoethanol and 20% glycerol using a NAP-5 column (GE Healthcare), flash frozen using liquid nitrogen, and stored at −80°C. Protein concentration was determined using Bradford reagent.

##### - Tubulin purification and labelling

Tubulin was purified from pig brains following the protocol described by Castoldi and Popov, based on cycles of polymerisation and depolymerisation and high-molarity buffer removal of associated proteins [37]. Fresh pig brain tissues were homogenized in 1 volume (1ml/g) of DB buffer (50mM MES pH 6.6, 1mM CaCl_2_) and centrifuged at 29,000g for 1h at 4°C. The supernatant was supplemented with 1 volume of high-molarity PIPES buffer (HMPB: 1M PIPES pH 6.8, 10mM MgCl_2_, 20 mM EGTA) and 1 volume of glycerol and raised to a final concentration of 1.5mM ATP and 0.5mM GTP. The mixture was then incubated for 1h at 37°C to induce tubulin polymerisation. After centrifugation at 150,000g for 30 min at 37°C, the pellet was resuspended in 0.3 volumes of cold DB buffer and incubated for 30 min at 4°C to depolymerize microtubules. After centrifugation at 4°C for 30 min at 120,000g, the supernatant containing free soluble tubulin was polymerized for a second cycle of 1h at 37°C after being supplemented as above with an equal volume of HMPB buffer, a volume of glycerol and with a final concentration of 0.5mM GTP and 1.5mM ATP. Polymerised tubulin was pelleted by centrifugation at 37°C for 30 min at 150,000g. The pellet was resuspended in 0.01 volumes of cold BRB80 buffer (80mM PIPES, 1mM EGTA, 1mM MgCl_2_, pH 6,8) and depolymerized for 1h at 4°C. The suspension was centrifuged for 30min at 4°C and 150,000g. The supernatant containing pure soluble tubulin was aliquoted, frozen in liquid nitrogen and stored at −80°C.

Labelling of tubulin with NHS-ester-ATTO 565 (ATTO-TEC) was performed following the protocol described by Hyman [38]. Tubulin was first polymerized in presence of 1mM GTP and 3.5mM MgCl_2_ and 25% glycerol (v/v) at 37°C for 1h. Microtubules were collected by centrifugation at 35°C for 40 min at 100,000g through a cushion of 0.1M HEPES pH 8.6, 1mM MgCl_2_, 1mM EGTA, 60% glycerol (v/v). The pellet containing microtubules was then resuspended in labelling buffer (0.1M HEPES pH 8.6, 1mM MgCl_2_, 1mM EGTA, 40% (v/v) glycerol). The succinimidyl ester-coupled fluorophore (dissolved at 50mM in DMSO) was then added to a final concentration of 5mM and incubated 20 min at 37°C. Labelled microtubules were centrifuged through a cushion of 60% (v/v) glycerol in BRB80 for 40min at 35°C at 150,000g. Microtubules were depolymerised for 30min at 4°C in 50mM K-Glutamate, 0.5mM MgCl_2_, pH 7.0 (KOH). Tubulin was recovered by centrifugation at 4°C for 20 min at 150,000g. The solution was brought to 80mM PIPES, 4mM MgCl_2_, 1mM GTP and another cycle of polymerisation and depolymerisation was performed. The final pellet was resuspended in cold BRB80, aliquoted, frozen in liquid nitrogen and stored at −80°C.

##### - Microtubule polymerisation

Tubulin and labelled tubulin (80μM total, with 7 to 20% labelling) were mixed in BRB80 with glycerol (80mM PIPES, 1mM EGTA, 1mM MgCl_2_, 25% glycerol, pH 6,8), and centrifuged to remove possible proteins aggregates or impurities (100,000g, 10min, 4°C). The supernatant was recovered in a microtube, 1mM GTP was added, and the mix was incubated 5min on ice, before incubation for polymerisation for 45min at 37°C. A volume of BRB80 supplemented with 40μM docetaxel (taxol analogue) was added to reach a concentration of 20μM docetaxel, and the mix was incubated for stabilization for 15 to 30 min (yielding more or less long MTs). The mix was centrifuged (100,000g, 10min, 30°C), the supernatant removed, and the pellet washed smoothly with 10μL BRB80D20 (BRB80 supplemented with 20μM docetaxel). BRB80D20 was added to the pellet, to resuspend the MTs at 47,8μM by pipetting smoothly after letting the pellet hydrate for 10min. MTs were stored at RT in the dark.

#### Buffers

We performed experiments in a dynein assay buffer (10mM K-PIPES, 50mM potassium acetate, 4mM MgSO_4_, 1mM EGTA, 10mM glucose, 25μM glucose oxydase, 6.4μM catalase, 1mM DTT, 0,4mg/mL casein, 40μM docetaxel, 1mM ATP, pH 7.0) supplemented with methylcellulose (MC) (Sigma). MC was previously prepared as a 2% solution as advised by the seller. Briefly, MC was added to half of the buffer volume heated to 80°C, agitated until particles are evenly dispersed, after which the remaining half of the buffer was added at 4°C. The mixture was then agitated at 4°C and MC dissolved when it attained the temperature at which it became water soluble. Glucose, DTT, casein, docetaxel and ATP were added after the solution cooled down to 4°C, and aliquots were stored at −20°C. Glucose oxydase, catalase and MTs were added just before experiments, to yield 1,18% or 1,60% MC.

#### Microscopy chambers

Chambers were made of two coverslips separated by one to three layers of parafilm. These coverslips were previously cleaned in ultrasound baths of Hellmanex 5%, KOH 2M, and absolute EtOH, or KOH 2M, demineralized H_2_0, and absolute EtOH (30min sonication in each solution, with H_2_0 rinsing in between each bath). Protein solutions were injected in the microscopy chambers with a pipet into chambers of 1.5 to 2mm in width, 50-440μm in height, and 0.5-2cm in length depending on experiments, resulting in a chamber volume of 3-8μL. The chambers were finally sealed with VALAP (an even mixture of vaseline, lanolin and paraffin).

#### Microscopy experiments

##### - Control experiments: gliding assay

A classical MT gliding assay was carried out to verify the functionality of the purified dynein, following the protocol described by Kon *et al*. [36]. Microscope chambers were constructed as described above, with one layer of parafilm between coverslips, a width of 2mm, an internal height of ~50μm, and a length of 2cm, resulting in a volume of ~6μL. The microscope chamber was covered with biotinamidocaproyl BSA, streptavidin, passivated with casein, (with buffer washes after each of these steps) and covered with 7nM biotinylated dynein. Finally, dynein assay buffer, supplemented with 400nM labelled and docetaxel-stabilized MTs, was flowed in the chamber. MTs were then free to meet dyneins at the surface and glide on them.

##### - Bead coating with dynein

Streptavidin beads were washed in the dynein assay buffer three times in Lobind microtubes (Sigma) (a wash included the mixing of beads in buffer, followed by a 2 minutes sonication, a centrifugation at 10,000g, 4min, 4°C, and the resuspension of the pellet in buffer). Beads were then incubated with biotinylated dynein for ~1-3h in the cold room with gentle agitation (incubation with more than 50 fold the beads binding capacity). Beads used were 0.5μm (Bangs laboratory), 1μm (Bangs laboratory or Dynabeads from ThermoFisher), 2.2μm (Polysciences), 2.8μm (ThermoFisher) and 3μm (Bangs laboratory). When beads were not already fluorescent, beads could be labelled with 0.001 fold the bead binding capacity with ATTO 565-biotin after the dynein coating step.

##### - Bead coating with kinesin

Streptavidin beads were washed as above, and then, either kinesin was incubated with 1μm beads (Bangs laboratory) for a direct adsorption on them, either beads were first incubated with biotin anti-GST, washed, and then incubated with kinesin. Each incubation consisted of ~1-3 hours of gentle agitation in a cold room, with more than 1000 fold the bead binding capacity, and each wash consisted of a centrifugation (10,000g, 4min, 4°C).

##### - Control experiments: bead gliding assay

Streptavidin beads were coated with dynein (as previously described). Microscope chambers were constructed as described above, with two layers of parafilm between the two coverslips, a width of 2mm, an internal height of ~200μm, and a length of 0,5cm, resulting in a volume of ~3μL. The microscope chamber was first covered with biotinamidocaproyl BSA and secondly with dynein-coated beads. Finally, dynein assay buffer supplemented with 400nM labelled and docetaxel-stabilized MTs, was flowed in the chamber.

##### - 3D motility assay experiment

The equivalent of approximately 0.005μL dynein-coated beads were centrifuged (10,000g, 6min, 20°C), re-suspended in ~3μL of 1.18% or 1.60% MC dynein assay buffer at 20°C supplemented with 12.5-25nM stabilized labelled MTs, and flowed in a hand-made chamber (width of 2mm, internal height of ~200μm, length of 0,5cm, resulting in a volume of ~3μL) which was then sealed with VALAP. We looked for events of a bead encountering a MT, focusing on MTs away from the surface (>10μm from the surface) and almost parallel to the surface. Data were acquired for ~1h.

##### - 3D motility assay experiments with optical trapping

The protocol was similar to the 3D motility assay described above, with the only difference that we did not looked for bead encountering MTs naturally, but instead captured beads with an home-made optical trap (IPG Ytterbium Fiber Laser (Model YLD-10-LP)) and placed them directly onto the MTs. This was particularly necessary to study the movement of two beads on a MT, or to construct a minimal aster and place several beads on it.

##### - Construction of an artificial minimal aster

Based on the protocol of the 3D motility assay with optical trapping, we increased the complexity of the *in vitro* reconstitution, and constructed a minimal artificial aster with 3μm dynein coated beads. A bead was placed on a MT, and as it walked on the MT, it revealed its polarity. The bead was detached from the MT by quickly pulling on it. This first step was repeated with 1 or 2 other MTs. The last MT handled was displaced thanks to the bead still present at its minus end, and with this bead, the other one or two MT precedently handled were captured by their minus ends. It resulted in a minimal aster, where two or three MT emanated from a 3μm bead. Two or three dynein coated beads could then be subsequently placed near the plus ends of the MTs of this minimal aster, to cause aster motion.

##### - Beads MSD measurement for viscosity assessment

To compute the viscosity of different media, we computed the Mean Square Displacement of beads of different sizes (1μm (Dynabeads,ThermoFisher), and 2.8μm (ThermoFisher)) in different medium. For this, beads were mixed in 1,18% or 1,60% MC dynein assay buffer, and frames were acquired every 1 or 2s for ~1min (Figure S2F). For each movies, we let the fluid equilibrate long enough, to ensure that flows were negligible, and did not affect the analysis.

##### - Hydrodynamic interaction experiment

Beads (1.9μm diameter, streptavidin coated) were labelled with ATTO 595-biotin, and mixed with stabilized labelled MTs in a viscous medium (1.18% MC dynein assay buffer). A single bead was captured with the optical trap, placed near a MT, and kept immobile there. The motorized stage was moved parallel to the MT, at velocities ranging from 1 to 45μm/s, resulting in a situation equivalent to the trapped bead being displaced near an immobile MT. The movement of the MT that may result from the bead displacement was measured to infer the hydrodynamic coupling between the two objects.

#### Acquisition

The microscopy chamber was placed on a Nikon EclipseTi or Nikon Eclipse TE2000-U inverted microscopes, using a 60x oil-immersion objective, with epifluorescence illumination. Microscopes were controlled through micromanager. The Nikon EclipseTi and Eclipse TE2000-U microscopes were illuminated with a Lumen Dynamics lamp (respectively X-Cite-Exacte and X-Cite-Series 120 Q) and movies were acquired by a Hamamatsu digital camera C11440 (respectively ORCA-Flash 4.0 and ORCA-Flash 4.0 LT plus)

### QUANTIFICATION AND STATISTICAL ANALYSIS

#### Specific experiments and analysis

Measurements were done manually using ImageJ data, and were treated using Microsoft Office Excel, Matlab and GraphPad Prism.

#### Classical gliding assay and bead gliding assay

In order to measure the MTs velocity on the dynein gliding assay, or the dynein-coated beads gliding assay, kymographs were constructed based on broken lines drawn along gliding MTs. Angles of MTs displacement were measured on the kymograph, and velocities were deducted from them.

#### 3D motility assay: motility of one or several beads on one MT

The analysis was carried on ImageJ: events were checked for persistent movements and constant velocity on kymographs, and distances travelled by the MT and the bead were measured on the movie (as shown in Figure S1). In addition to brownian motion, moderate local fluid flows in the microchamber added some noise to our measurements. MT length was measured as the length of a broken line following the MT from its minus end to its plus end, on one frame of the movie, or, when the MT was not in perfect focus trough the time-lapse, on a small z stack done at the end of the movie. MT bundling was assessed by comparing the MT fluorescence with the one of single MTs in the same field of view.

#### 3D motility assay: motility of several beads on a minimal aster

The movements of beads were tracked on ImageJ, following the bead with a circle and measuring its centroid, using a sub-sampling of 1/100^th^ of the pixel size to increase the measurement precision. Velocities of walking beads and aster center bead could then be deduced from these positions.

#### MSD measurement for viscosity assessment

We selected beads in focus to be tracked, and checked that they were single beads (and not aggregates) with a z stack acquisition at the end of the movie. Beads centers were tracked with the ImageJ manual tracking tool. Mean Square Displacements (MSD) were then plotted as a function of the time, and the diffusion coefficient D was deduced from the slope of the fit. This allowed to compute a diffusion coefficient, from the relation 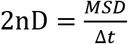, with n the number of dimensions (here n = 2 as we are measuring the MSD in 2D). The viscosity felt by the beads could then be calculated, using the Stokes–Einstein equation, for diffusion of spherical particles in a liquid at low Reynolds number: 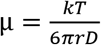, with *k* the Boltzmann constant, *T* the temperature and *r* the beads radius.

#### Relationship between speeds and dimensions of MT and beads

##### For one bead on one MT

Let us first consider a single motor-coated bead moving along a single MT, in solution, away from all surfaces. Neglecting drift and diffusion, all movements occur along the axis materialized by the MT, which we arbitrarily choose to orient towards the MT minus end. In the laboratory frame of reference, the bead and the MT move with velocities V_bead_ and V_MT_, respectively. For the isolated system (bead + MT), the force balance yields: γ_MT_ v_MT_ + γ_bead_ v_bead_=0, where γ_MT_ and γ_bead_ are the MT longitudinal drag coefficient, and the bead drag coefficient, respectively.

Based on Stoke’s law, the drag coefficient of the bead is expected to scale with the bead radius, R_bead_. Within our range of MT lengths, we can reasonably approximate the longitudinal drag coefficient of the MT as a linear function of the MT length, L_MT_ [2, 25]. We can thus write γ_bead_ = μ α_bead_ R_bead_ and γ_MT_ = μ α_MT_ L_MT_, where α_bead_ and α_MT_ are geometrical constants, and μ is the viscosity of the medium. As a consequence, the speed ratio -V_bead_/V_MT_ = α_MT_L_MT_ / α_bead_R_bead_ is expected to scale with the MT-length-to-bead-radius ratio, independently of the viscosity of the medium.

In order to assess how efficiently the activity of the motor is converted into MT movement, it is convenient to compute the MT-to-motor speed ratio -V_MT_/v*_motor_, where v*_motor_ is the speed at which the motor transports the bead along the MT, and which can be written v*_motor_=V_bead_-V_MT_. Based on the previous equations, this “efficiency” ratio can be written as: -V_MT_/v*_motor_ = 1/(1+ α_MT_L_MT_ / α_bead_R_bead_), and is comprised between 0 and 1.

Fitting the experimental plot of -v_MT_/v*_motor_ versus L_MT_/R_bead_ (Figure 2E) with this equation provides an estimation of α_MT_/α_bead_, which is the only free parameter of the fit. We thus find this number to be 0.020, which is approximately 2.3-fold smaller than what can be computed theoretically for a Newtonian medium, based on Stoke’s equation and on our linear approximation of the MT longitudinal drag coefficient (Figure S3A). This difference may be explained by the fact thatthe buffer supplemented with MC may behave as a non-Newtonian fluid, and thus exhibit significant shear-thinning effects [29]. In addition, our theoretical computation also neglects potential complications such as the hydrodynamic coupling between the bead and the MT, and the rotation of the bead as it moves along the MT.

##### For two beads moving along a single MT

Considering that the drag forces of two motile beads are independent and additive, the force balance yields γ_MT_ V_MT_ + γ_bead1_ V_bead1_ + γ_bead2_ V_bead2_ =0, and the computation of the ratio - V_MT_/v*_motor_ then leads to:

-V_MT_/<v*_motor_> = 1/(1+ α_MT_L_MT_ / 2α_bead_R_bead_), where <v*_motor_>=(v*_motor1_+v*_motor2_)/2 is the average motor speed (Figure 3C).

##### For multiple beads moving on a minimal aster

We now consider the movement of N motor-coated beads on a set of MTs connected by their minus ends to a central bead. Movements are observed in the focal plane of the microscope, in two dimensions. The minimal aster is thus composed of the central bead and the MTs, and has a non-isotropic drag coefficient. Balancing the drag forces yields, projected on a x-axis parallel to the two MTs in the minimal aster:

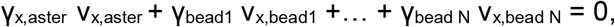

where γ_x,aster_ is the drag coefficient of the minimal aster for movements along the x-axis.

Thus v_x,aster_ should scale with −(v_x,bead1_ +… + v_x,bead N_). (Figure 4B).

#### Hydrodynamic interaction experiment (Figure S4)

An optically trapped bead was displaced near a freely floating MT, by keeping the trapped bead in a fixed position, in the frame of reference of the microscope, while moving the entire chamber thanks to a motorized stage. A free bead, away from the MT and the trapped bead, was used to monitor the movement of the stage, in the frame of reference of the microscope. All movements were then determined in the frame of reference of the stage, by using the free bead as a reference.

The velocity of the trapped bead was thus measured, in the frame of reference of the stage, and so was the velocity of the MT, resulting from the hydrodynamic coupling with the trapped bead displaced parallel to it. The MT-to-bead speed ratio can be plotted as a function of the MT distance from the surface of trapped bead, and data can be fitted as a power series (Figure S4C).

### KEY RESOURCES TABLE

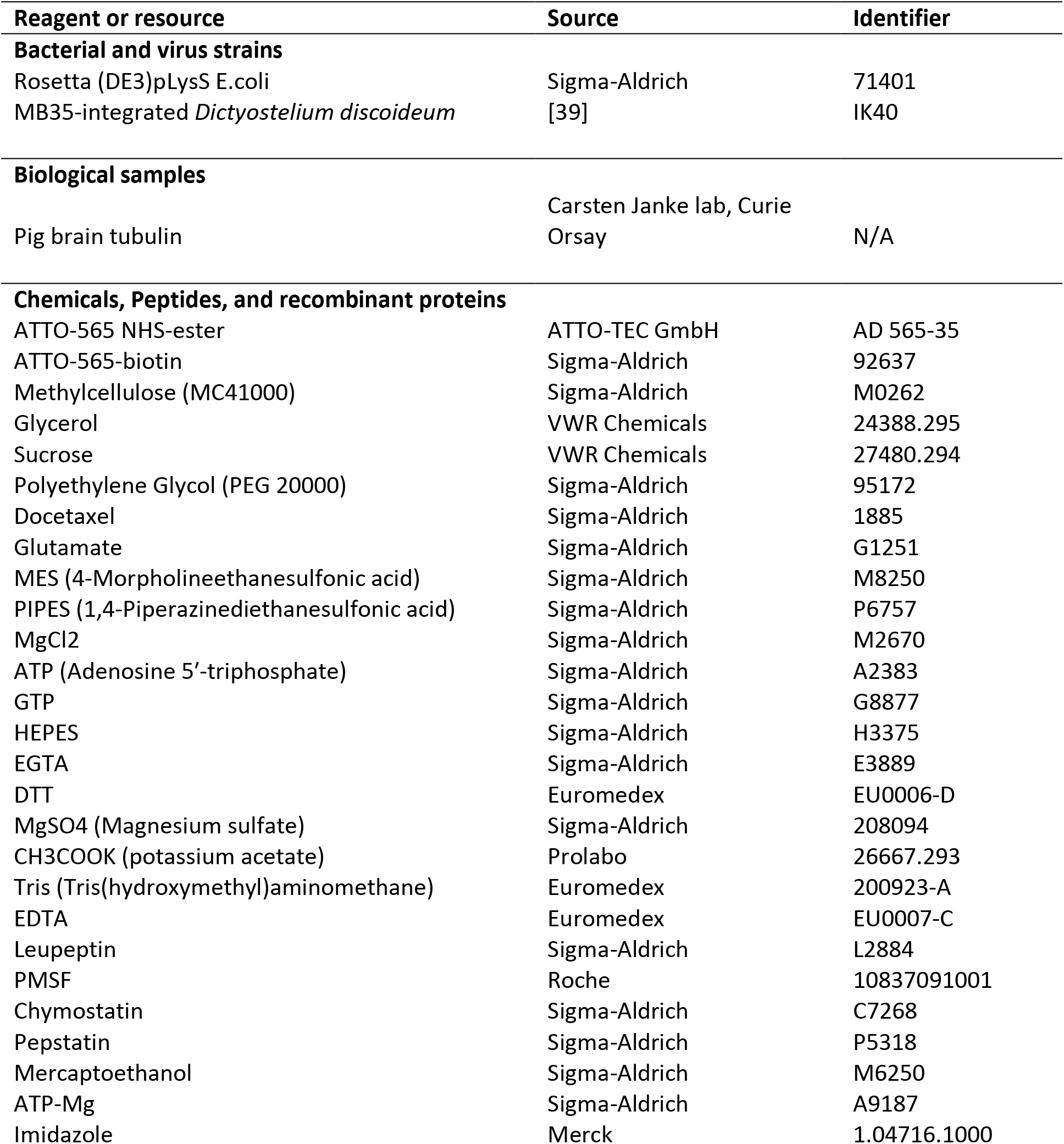
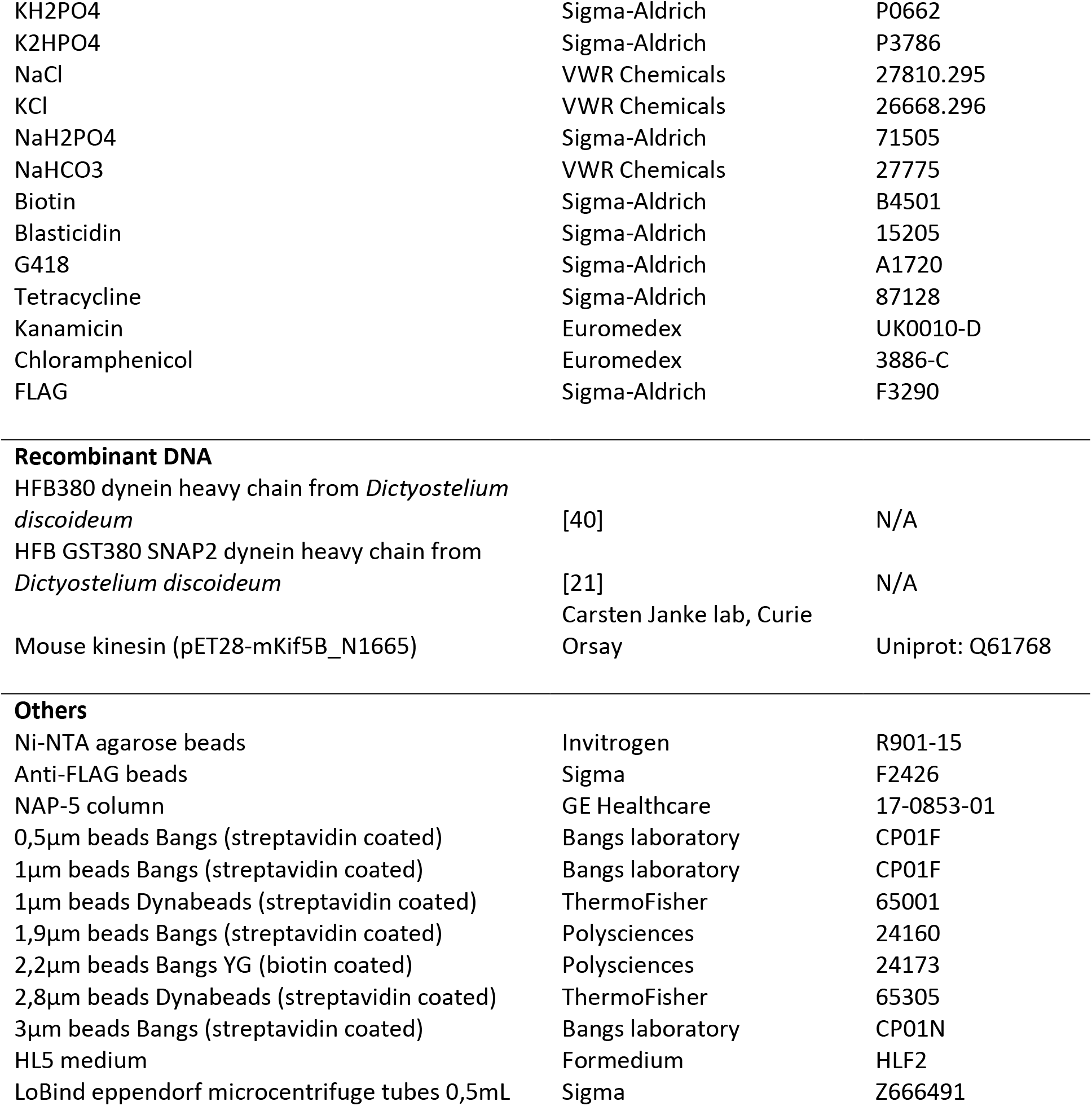

| Reagent or resource | Source | Identifier |
| --- | --- | --- |
| **Bacterial and virus strains** |  |  |
| Rosetta (DE3)pLysS E.coli | Sigma-Aldrich | 71401 |
| MB35-integrated *Dictyostelium discoideum* | [39] | IK40 |
| **Biological samples** |  |  |
| Pig brain tubulin | Carsten Janke lab, Curie Orsay | N/A |
| **Chemicals, Peptides, and recombinant proteins** |  |  |
| ATTO-565 NHS-ester | ATTO-TEC GmbH | AD 565-35 |
| ATTO-565-biotin | Sigma-Aldrich | 92637 |
| Methylcellulose (MC41000) | Sigma-Aldrich | M0262 |
| Glycerol | VWR Chemicals | 24388.295 |
| Sucrose | VWR Chemicals | 27480.294 |
| Polyethylene Glycol (PEG 20000) | Sigma-Aldrich | 95172 |
| Docetaxel | Sigma-Aldrich | 1885 |
| Glutamate | Sigma-Aldrich | G1251 |
| MES (4-Morpholineethanesulfonic acid) | Sigma-Aldrich | M8250 |
| PIPES (1,4-Piperazinediethanesulfonic acid) | Sigma-Aldrich | P6757 |
| MgCl<sub>2</sub> | Sigma-Aldrich | M2670 |
| ATP (Adenosine 5'-triphosphate) | Sigma-Aldrich | A2383 |
| GTP | Sigma-Aldrich | G8877 |
| HEPES | Sigma-Aldrich | H3375 |
| EGTA | Sigma-Aldrich | E3889 |
| DTT | Euromedex | EU0006-D |
| MgSO<sub>4</sub> (Magnesium sulfate) | Sigma-Aldrich | 208094 |
| CH<sub>3</sub>COOK (potassium acetate) | Prolabo | 26667.293 |
| Tris (Tris(hydroxymethyl)aminomethane) | Euromedex | 200923-A |
| EDTA | Euromedex | EU0007-C |
| Leupeptin | Sigma-Aldrich | L2884 |
| PMSF | Roche | 10837091001 |
| Chymostatin | Sigma-Aldrich | C7268 |
| Pepstatin | Sigma-Aldrich | P5318 |
| Mercaptoethanol | Sigma-Aldrich | M6250 |
| ATP-Mg | Sigma-Aldrich | A9187 |
| Imidazole | Merck | 1.04716.1000 |
| KH<sub>2</sub>PO<sub>4</sub> | Sigma-Aldrich | P0662 |
| K<sub>2</sub>HPO<sub>4</sub> | Sigma-Aldrich | P3786 |
| NaCl | VWR Chemicals | 27810.295 |
| KCl | VWR Chemicals | 26668.296 |
| NaH<sub>2</sub>PO<sub>4</sub> | Sigma-Aldrich | 71505 |
| NaHCO<sub>3</sub> | VWR Chemicals | 27775 |
| Biotin | Sigma-Aldrich | B4501 |
| Blasticidin | Sigma-Aldrich | 15205 |
| G418 | Sigma-Aldrich | A1720 |
| Tetracycline | Sigma-Aldrich | 87128 |
| Kanamycin | Euromedex | UK0010-D |
| Chloramphenicol | Euromedex | 3886-C |
| FLAG | Sigma-Aldrich | F3290 |
| **Recombinant DNA** |  |  |
| HFB380 dynein heavy chain from *Dictyostelium discoideum* | [40] | N/A |
| HFB GST380 SNAP2 dynein heavy chain from *Dictyostelium discoideum* | [21] | N/A |
| Mouse kinesin (pET28-mKif5B_N1665) | Carsten Janke lab, Curie Orsay | Uniprot: Q61768 |
| **Others** |  |  |
| Ni-NTA agarose beads | Invitrogen | R901-15 |
| Anti-FLAG beads | Sigma | F2426 |
| NAP-5 column | GE Healthcare | 17-0853-01 |
| 0,5µm beads Bangs (streptavidin coated) | Bangs laboratory | CP01F |
| 1µm beads Bangs (streptavidin coated) | Bangs laboratory | CP01F |
| 1µm beads Dynabeads (streptavidin coated) | ThermoFisher | 65001 |
| 1,9µm beads Bangs (streptavidin coated) | Polysciences | 24160 |
| 2,2µm beads Bangs YG (biotin coated) | Polysciences | 24173 |
| 2,8µm beads Dynabeads (streptavidin coated) | ThermoFisher | 65305 |
| 3µm beads Bangs (streptavidin coated) | Bangs laboratory | CP01N |
| HL5 medium | Formedium | HLF2 |
| LoBind eppendorf microcentrifuge tubes 0,5mL | Sigma | Z666491 |

