## Supplemental data for "In Vitro Reconstitution of Dynein Force Exertion in Bulk Cytoplasm"

### SUPPLEMENTAL FIGURES

A

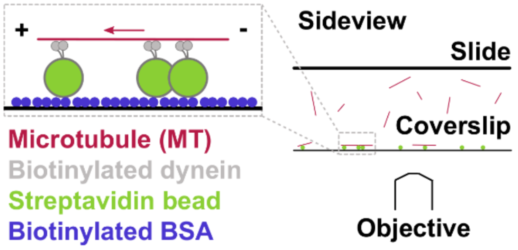


B

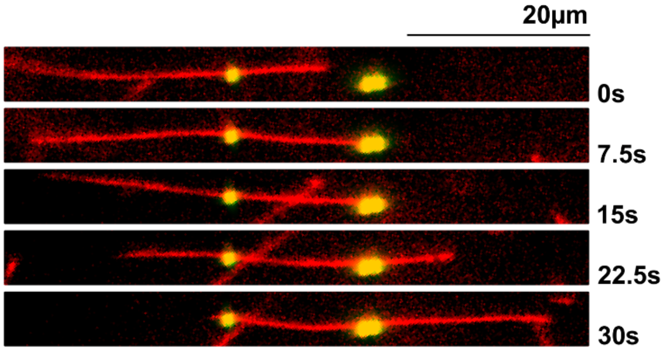


C

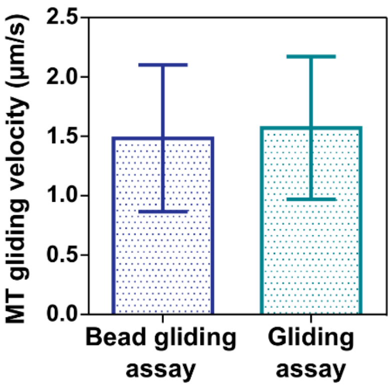


D

| Viscous agents | Effect on dynein activity | MT bundling |
|---|---|---|
| Sucrose | Inhibition [S1]<br>sucrose 20%, i.e. ~2cP → 1.13μm/s ± 0.40<br>sucrose 47%, i.e. ~11cP → 0.06μm/s ± 0.01 | • sucrose 70% - [tubulin] 400nM<br>→ Single MTs |
| Glycerol | Inhibition [S1, S2] | • glycerol 68% - [tubulin] 400nM<br>→ Single MTs |
| Polyethylene Glycol 20000 ($PEG_{20000}$) | Inhibition<br>when [$PEG_{20000}$] > 2% [S1, S3, S4] | • $PEG_{20000}$ 30% - [tubulin] 400nM<br>→ Single MTs or ~5-MT bundles |
| Methylcellulose 41000 ($MC_{41000}$) | No strong inhibition<br>$MC_{41000}$ 1.41%, i.e. ~100cP → 0.39μm/s ±0.21 | • $MC_{41000}$ 2% - [tubulin] 400nM<br>→ Single MTs or ~4-MT bundles<br>• $MC_{41000}$ 2% - [tubulin] 10nM<br>→ Single MTs or ~2-MT bundles |

E

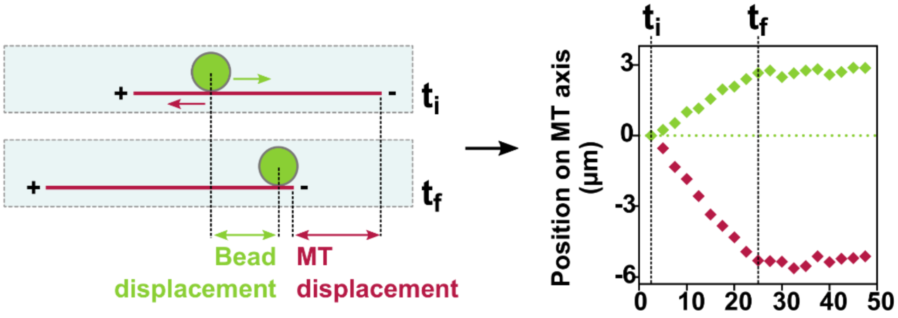

**<u>Figure S1:</u> Control experiments for the 3D motility assay: dynein coating functionality on beads, and MTs and dynein behavior in different viscous agents. Related to Figure 1.**

(A) Sketch representing the bead surface gliding assay. Biotinamidocaproyl BSA is flowed in a microscopy chamber. Streptavidin beads coated with biotinylated dynein are then incubated in the chamber, and finally stabilized MTs are flowed in and can freely glide on dynein-coated beads anchored to the surface. (see STARMethods).

(B) Time-lapse of a stabilized MT gliding on three anchored dynein-coated beads, in a typical bead gliding assay experiment.

(C) Average velocity of MTs gliding on a dynein-coated beads gliding assay (Dynabeads, 1µm diameter) or a conventional dynein gliding assay, both in dynein assay buffer. N=42 MTs (bead gliding assay) and 29 MTs (gliding assay). Error bars represent standard deviation.

(D) Table showing all viscous agents tried: sucrose, glycerol, PEG (molecular weight 20000 g/mol) and MC (molecular weight 41000 g/mol), and their potential inhibition of dynein ATPase activity and motility (from our gliding assays MTs speeds, or from the literature) and their tendency to generate MT bundles when supplemented to our dynein assay buffer.

(E) Sketch showing the measurements done to analyse the 3D motility assay and an example of the choice for $t_i$ and $t_f$, encompassing a constant bead and MT movement.

**A**

| Bead diameter (µm) (1µm Dynabeads) | |
|---|---|
| Provided | Measured |
| 1.05 ± 0.032 | 0.96 ± 0.032 |

**B** Dynein velocity (3D motility assay, 1µm beads)

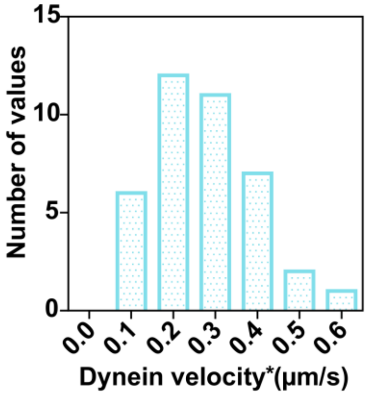


**C**

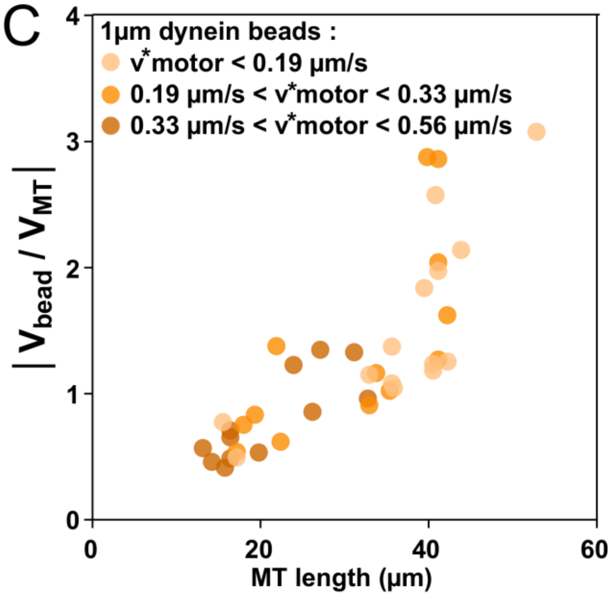


**D** Kinesin and dynein velocity (3D motility assay)

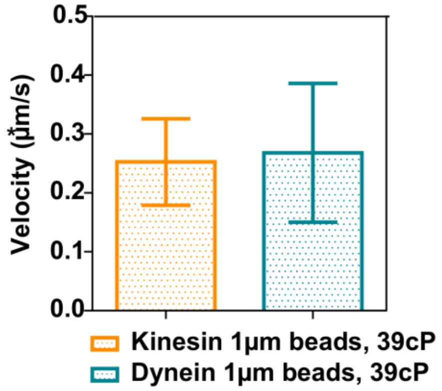


**E**

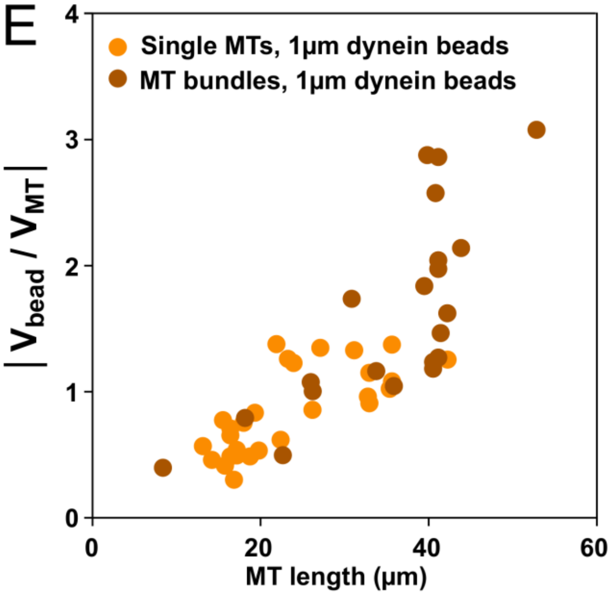


**F** Example of diffusion coeficient D determination from MSD measures

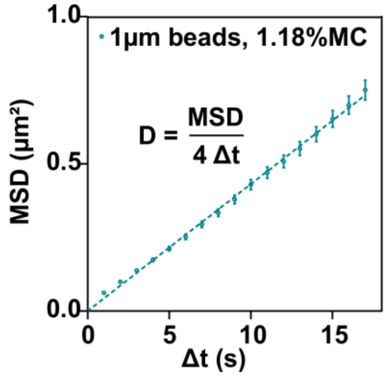


**G** Viscosity determined from measured diffusion coeficient

| | Viscosity from MSD (cP) | |
|---|---|---|
| | 1.18% MC | 1.60%MC |
| 1µm beads | 37.9 ± 0.2 | 123.5 ± 0.1 |
| 2.8µm beads | 40.7 ± 0.6 | \ |

**<u>Figure S2:</u> Experimental determination of medium viscosity, comparison of kinesin and dynein coated beads velocities on MTs, and independence of the 3D motility assay behaviour on motor velocity or the presence of thin MT bundles. Related to Figure 1 and 2.**

(A) Bead size control of the 1µm beads (Dynabeads). We compare the specifications provided by the manufacturer, and the Feret diameter measured with the “analyze particle” plugin in ImageJ, for beads sedimented on the surface of a coverslip and observed in transmitted light.

(B) Histogram of dynein velocities, $v^*_{motor}$, on the MTs in the 3D motility assay experiment, in 39cP MC dynein assay buffer, for 1µm beads (Bangs laboratory).

(C) Bead-to-MT speed ratio plotted as a function of MT length, for 1µm diameter beads (Bangs laboratory), for different ranges of motor velocity, in a 39cP MC viscous medium.

(D) Average velocity of the dynein and kinesin motors in the 3D motility assay experiment (1µm diameter beads from Bangs laboratory, in a 39cP MC dynein assay buffer). N=39 beads (dynein) and 11 beads (kinesin). Error bars represent standard deviation.

(E) Bead-to-MT speed ratio, as a function of the MT length, for 1µm diameter beads (Bangs laboratory and Dynabeads), for single MTs and small MT bundles (2- and 3-MT bundles), in a 39cP and 124cP MC viscous medium.

(F) Example of the determination of the diffusion coefficient of 1µm beads (Dynabeads) in the 1.18% MC dynein assay buffer through MSD measurements. Error bars represent standard deviations.

(G) Table of viscosity assessment through diffusion coefficient determination for 1µm and 2.8µm beads (Dynabeads), and 1.18% and 1.60% methylcellulose (MC) dynein assay buffers (thus elsewhere named 39cP and 124cP MC viscous buffers).

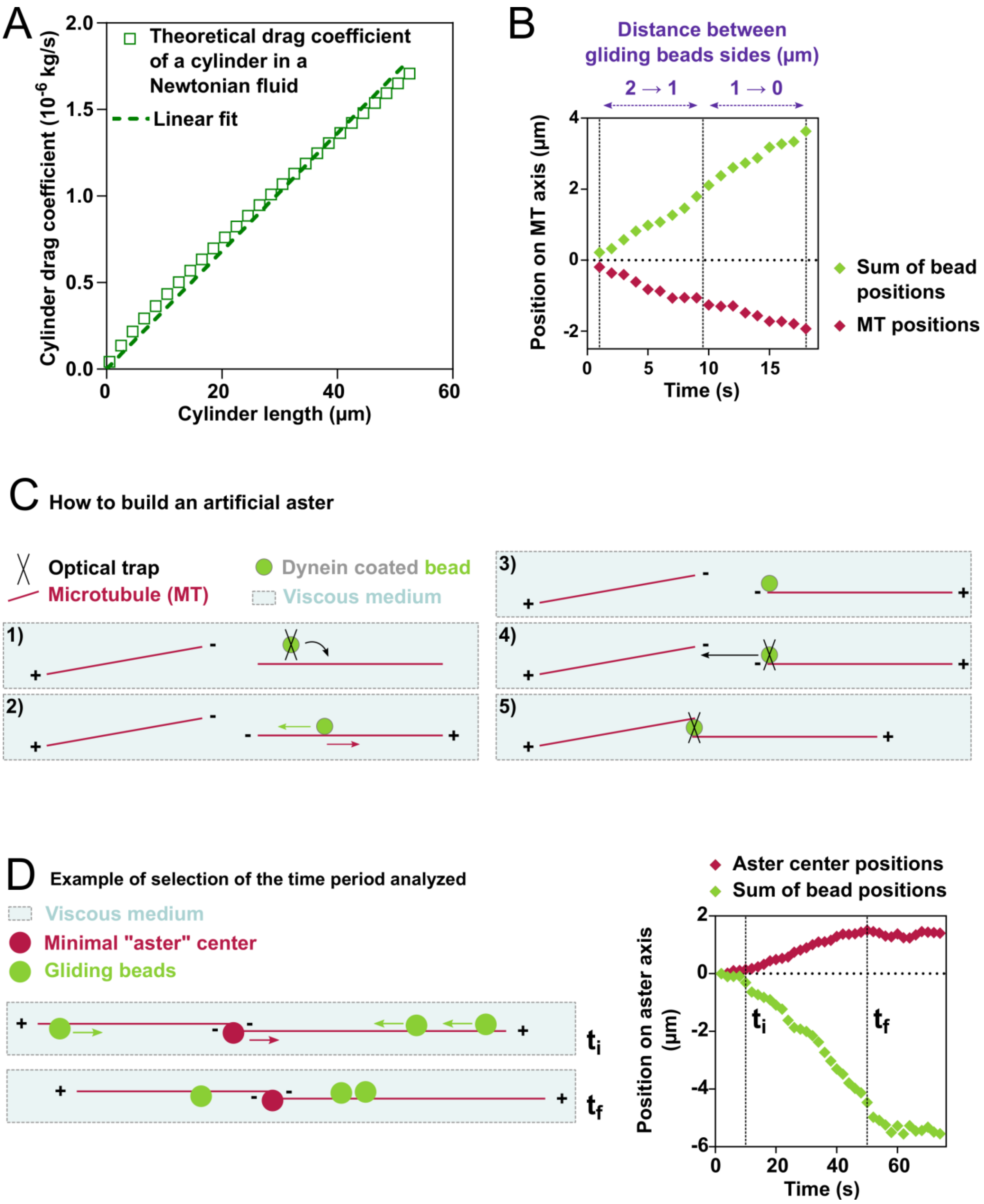
A
Theoretical drag coefficient of a cylinder in a Newtonian fluid
Linear fit
Cylinder drag coefficient (10^-6 kg/s)
Cylinder length (µm)
B
Distance between gliding beads sides (µm)
2 → 1
1 → 0
Position on MT axis (µm)
Time (s)
Sum of bead positions
MT positions
C How to build an artificial aster
Optical trap
Microtubule (MT)
Dynein coated bead
Viscous medium
1)
2)
3)
4)
5)
D Example of selection of the time period analyzed
Viscous medium
Minimal "aster" center
Gliding beads
ti
tf
Aster center positions
Sum of bead positions
Position on aster axis (µm)
Time (s)

**Figure S3: Theoretical longitudinal drag coefficient of a cylinder, and method to build a minimal artificial aster. Related to Figure 3 and 4.**

(A) Linear approximation of a cylinder's longitudinal drag coefficient, computed for a 25 µm diameter cylinder in a newtonian fluid, with viscosity 39 cP, with a theoretical value of $\frac{2\pi\, \upsilon\ L_{MT}}{\ln\left(\frac{L_{MT}}{2r_{MT}}\right)+\gamma_{\|}}$, with $L_{MT}$ and $r_{MT}$the length and radius of the cylinder, $\upsilon$ the viscosity and $\gamma_{||}$ the end correction term.

(B) Beads sum positions and MT positions plotted as a function of time, in a configuration where two 1µm dynein coated beads glide on a MT. The MT velocity does not change significantly even when beads are getting closer to each other.

(C) Sketch explaining the steps of construction of a minimal artificial aster. A bead is first placed on a MT, and as it walks on the MT it reveals its polarity (1-2). The bead is then detached from the MT by sharply pulling on it. This first step is repeated with another MT (3-4). This last handled MT is displaced thanks to the bead still present at its minus end, and with this bead, the other one MT precedently handled is captured by its minus ends (5).

(D) Example showing how tug-of-war events implicating multiple beads are analyzed.

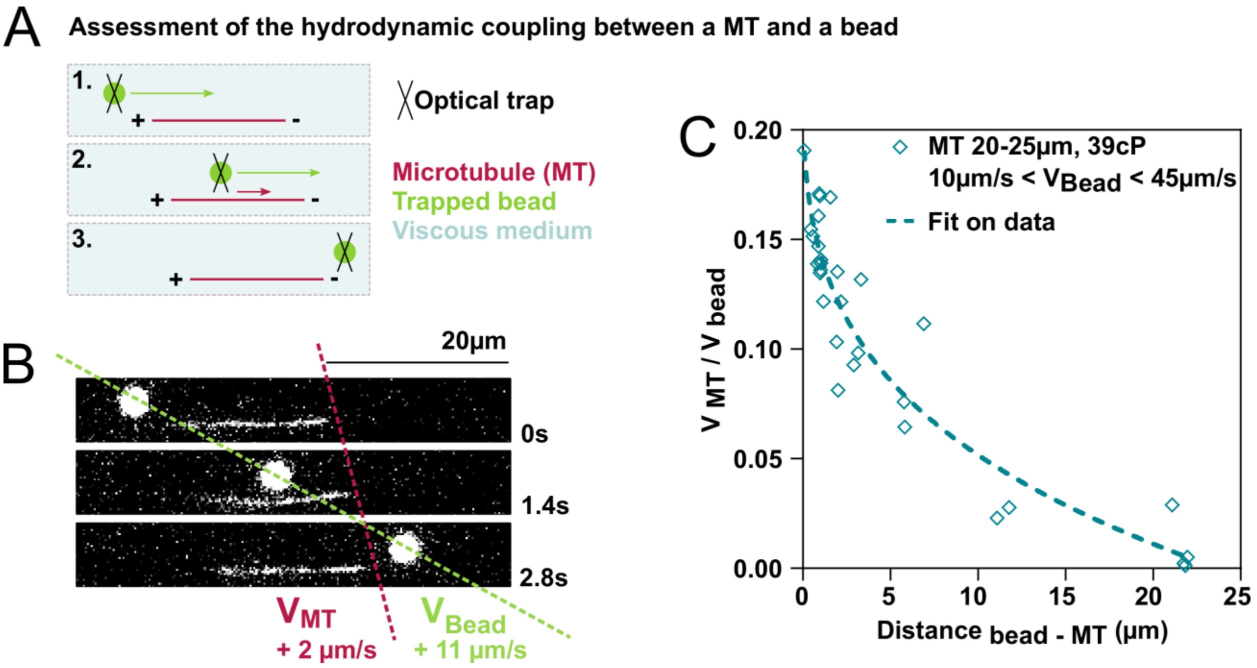
A
Assessment of the hydrodynamic coupling between a MT and a bead
1.
2.
3.
Optical trap
Microtubule (MT)
Trapped bead
Viscous medium
B
20µm
0s
1.4s
2.8s
$V_{MT}$
+ 2 µm/s
$V_{Bead}$
+ 11 µm/s
C
MT 20-25µm, 39cP
10µm/s < $V_{Bead}$ < 45µm/s
Fit on data
$V_{MT}$ / $V_{bead}$
Distance bead - MT (µm)
0.00
0.05
0.10
0.15
0.20
0
5
10
15
20
25

**<u>Figure S4:</u> Hydrodynamic coupling between a MT and bead moving in its vicinity. Related to Figure 1 to 4.**

(A) Sketch of the hydrodynamic interaction assay. A bead is moved parallel to a nearby MT, and the subsequent MT movement can be measured, as a function of the MT length and MT distance to the bead.

(B) Time-lapse of an example of the hydrodynamic interaction assay. One can note how the MT is being displaced through the hydrodynamic coupling with the bead displaced parallel to this MT.

(C) MT-to-bead speed ratio plotted as a function of the distance from the side of the bead to the MT, for MT between 20 and 25µm, bead velocity from 10 to 45µm/s and beads of 1.9µm diameter (Polysciences), in a 1.18% MC viscous medium (39cP viscosity).

**MOVIE LEGENDS**

**Movie S1: 3D motility assay**

**Part 1: a small bead walks on a long MT**

Acquired in epifluorescence (stabilized MTs, 20% tubulin ATTO-565, in red; green fluorescent beads, in green), away from the coverslip surface, at 1 frame / 2s. Scale bar 20µm. The bead diameter is 0.5µm, the MT length is 36µm.

**Part 2: a large bead walks on a short MT**

Acquired in epifluorescence (stabilized MTs, 20% tubulin ATTO-565; ATTO-565 bead, both in white), away from the coverslip surface, at 1 frame / 3s. Scale bar 20µm. The bead diameter is 3µm, the MT length is 24µm.

**Movie S2: How to place 2 beads on a MT with an optical trap**

Acquired in epifluorescence (stabilized MTs, 15% tubulin ATTO-565; beads auto fluorescent in red, both in white), away from the coverslip surface.

**Movie S3: 3D motility assay: two beads walk on a MT**

Acquired in epifluorescence (stabilized MTs, 7% tubulin ATTO-565; beads auto fluorescent in red, both in white), away from the coverslip surface, at 1 frame / 1s. Scale bar 20µm. Beads diameter is 1µm, MT length is 23µm.

**Movie S4: Tug-of-war**

**Part 1: three motile beads walk on a two MTs minimal aster**

Acquired in epifluorescence (stabilized MTs, 20% tubulin ATTO-565, ATTO-565 beads, both in white), away from the coverslip surface, at 1 frame / 2s. Scale bar 20µm. Beads diameter is 3µm. The minimal aster is composed of two MTs attached to a central bead, and three motile beads are placed on the MTs, before acquiring the movie.

**Part 2: three motile beads walk on a three MTs minimal aster**

Acquired in epifluorescence (stabilized MTs, 20% tubulin ATTO-565, ATTO-565 beads, both in white), away from the coverslip surface, at 1 frame / 2s. Scale bar 20µm. Beads diameter is 3µm. The minimal aster is composed of three MTs attached to a central bead, and three motile beads are placed on the MTs, before acquiring the movie.
